# The phenotypic nonspecificity of cell-to-cell signalling in *Drosophila melanogaster*

**DOI:** 10.64898/2026.05.19.726339

**Authors:** Anthony Percival-Smith, Cooper Brabrook

## Abstract

An expectation of a hypothesis that proposes cell-to-cell signalling pathways are redundant due to the redundancy of pathway terminal transcription factors (TFs) was tested by screening 35 signalling ligands (SLs) for rescue of a decapentaplegic (dpp) hypomorphic wing growth phenotype. The screen identified three examples of partial rescue: Hedgehog (HH), Semphorin 1a (SEMA1A) and Wnt ortholog 2 (WNT2). HH overexpression with *dppGAL4* may increase the expression of DPP activity from the hypomorphic *dpp* alleles. However, SEMA1A and WNT2 did not phenocopy ectopic expression of HH or DPP and neither SEMA1A nor WNT2 were required for wing growth suggesting substitution of DPP for partial restoration of wing growth. The WNT2 rescue was dependent on the Frizzled 4 (FZ4) WNT receptor excluding the possibility that WNT2 weakly binds the DPP receptor. Although examples of phenotypic nonspecificity of SL function were identified, this is an expectation, and not direct proof, of the hypothesis of TF redundancy.

**Screen Report Summary:** An expectation of a hypothesis proposing that cell-to-cell signalling pathways are redundant due to the redundancy of the pathway terminal transcription factors was tested by screening for replacement of one signalling ligand (DPP; SLa) with another SLb for wing growth. Three non-DPP SLs were identified in the screen of 35SLs: HH, SEMA1A and WNT2. Genetic analysis of *Sema1a* and *Wnt2* suggests functional complementation of *dpp* for wing growth suggesting that SEMA1A and WNT2 partially replace DPP for wing growth. Therefore, an expectation of the hypothesis is met.

## Introduction

Cell-to-cell signalling is essential for the development and physiology of multicellular and single-celled organisms. One general sequence of information flow common to many signalling pathways is the specific interaction of a ligand with a transmembrane receptor on the surface of the cell followed by the activation of an intracellular pathway that leads to changes in the activity of a pathway terminal TF(s), thereby altering gene expression in the cell. The protein encoded by the Drosophila *dpp* gene is a cell-to-cell signalling ligand of the Tumor Growth Factor β (TGF-β) ligand family and the Drosophila ortholog of vertebrate bone morphogenetic proteins (Padgett et al., 1987). DPP is required for many different developmental processes of which a limited list includes embryonic dorsal ventral axis formation, axis formation of the developing adult leg, wing growth and patterning, eye formation and gonad development (Chanut and Heberlein 1997; Posakony et al., 1990; Xie and Spradling 1998). In both embryonic dorsal ventral axis formation and wing morphogenesis, DPP is proposed to function as a concentration-dependent morphogen (Ferguson and Anderson 1992; Nellen et al., 1996). However, subsequent studies of the role of DPP spreading during wing morphogenesis do not support DPP acting as a simple concentration-dependent morphogen in wing pattern formation (Akiyama and Gibson, 2015; Harmansa et al., 2015; Matsuda and Affolter, 2023). The protein complex consisting of the DPP ligand bound to the Thickvein Punt transmembrane receptor phosphorylates the transcription factor Mothers against dpp (MAD), and the phosphorylated MAD forms a protein complex with another TF Medea (MED) that together enter the nucleus to alter gene expression (Padgett et al., 1998).

The functional redundancy of TFs results in the phenomenon of phenotypic nonspecificity, which is the substitution of one TFa with that of another TFb for the rescue of the TFa phenotype (Percival-Smith 2017; Percival-Smith et al., 2023). Combining the observations that signal transduction pathways affect TF activity and that TF activity is redundant suggests the possibility that cell-to-cell signalling pathways are also phenotypically nonspecific. Phenotypic nonspecificity of cell-to-cell signalling would be the substitution of one signalling factor, SLa by another unrelated SLb for rescue of the SLa phenotype. This recue is the hypothetical result of the TFs affected by the presence/absence of the SLs a or b being functionally redundant. This expectation of the TF redundancy hypothesis was assessed by screening a set of Drosophila signalling factors expressed with a *dppGAL4* driver for wing growth rescue of a *dpp* hemizygous, hypomorphic mutant combination (Spencer et al., 1982).

## Materials and Methods

### Drosophila husbandry and the screen

Flies were maintained at 24 °C and 60% humidity. The flies were reared in 20ml vials containing corn meal media [1% (w/v) Drosophila grade agar, 6% (w/v) sucrose, 10% (w/v) cornmeal, 1.5% (w/v) yeast, and 0.375% (w/v) 2-methyl hydroxybenzoate]. The genotypes of the stocks used in this study are given in Table S1. The stocks were where used in standard Drosophila crosses to assemble the specific genotypes used in this study. Flybase was first screened for genes that express secreted signalling ligands using the GO term “receptor ligand activity”, and this subset of genes was then screened for available stocks at the Bloomington and FlyORF Drosophila stock centers with UAS insertions expressing the various secreted ligands on the third chromosome (Öztürk-Çolak 2024). The UAS fly stocks were used to create the common genotype *y w; dpp*^*d5*^*/CyO; P or M{UASX, w*^+^*}* where X is the ligand expressed. For the screen, virgins of this common genotype were crossed with *y w; dpp*^*d6*^*/L; P{dppGAL4, w*^+^*}*/+ males, which were generated by crossing virgin *y w; CyO/L; P{dppGAL4, w*^+^*}/TM6B* females with *dpp*^*d6*^*/CyO* males. The *y w; dpp*^*d5*^*/dpp*^*d6*^; *P or M{UASX, w*^+^*}/ P{dppGAL4, w*^+^ progeny were screened for rescue of the dpp hypomorphic wing phenotype (*P{dppGAL4, w*^+^*}* progeny have dark red eyes).

### Imaging rescue and quantification of wing area

For imaging the rescued wings with scanning electron microscopy, flies were first critical point dried and then sputter coated. The images were collected using a scanning electron microscope. For wing area measurements, adult wings were plucked and placed in 70% ethanol. The wings were mounted on slides in Hoyer’s mountant (Wieschaus and Nusslein-Volhard, 1986). The wing area was collected on a Zeiss AxioImager Z1 microscope using Zen imaging software (Carl Zeiss Canada Inc.). DPP expression was knocked down in the progeny of 38418 X 25782 (TableS1).

### Analysis of the requirement of the Frizzled 4 receptor

To assess whether the Frizzled 4 receptor (FZ4) was required for rescue of the dpp phenotype by expression of WNT2, the *fz4*^*3-1*^ null mutant allele (McElwain et al., 2011) was used to generate *y w fz4*^*3-1*^; *dpp*^*d5*^*/CyO; M{UASWnt2, w*^+^*}* virgin females that were crossed with *y w; dpp*^*d6*^*/CyO; P{dppGAL4, w*^+^*}*/+ males. The *y w fz4*^*3-1*^; *dpp*^*d5*^*/ dpp*^*d6*^; *M{UASWnt2, w*^+^*}/; P{dppGAL4, w*^+^*}* male progeny were assessed for rescue.

### RNA interference

Male flies from stock 38418 were crossed with 24645 virgin females, and male *P{UASDcr-2}, w*^*1118*^; *P{nubGAL4, w*^+^*}/ TM3 Sb* progeny collected and crossed with 29554, 30483, 34320 and 67845 virgin females (Table S1; Ni et al., 2009). For measuring wing area, the wings were plucked from female flies that had neither third nor second chromosome balancers. For quantification of *PLEXA* mRNA, 38418 males homozygous for the *nubGAL4* driver were crossed with either 30483 or 67845 homozygous for *UAS*{*RNAi}* and wing imaginal discs dissected from late third instar larval progeny.

### Quantitative PCR

Male and female larvae were distinguished by the characteristic sexually dimorphic size of their gonads. 10-20 wing discs were dissected from late third instar larvae and total RNA extracted with a RNeasy mini kit (QIAGEN). The RNA was reversed transcribed with SuperScript IV (Thermofisher). The qPCR was performed with the GB-Amp™ PowerGreen™ qPCR mix (Genebio Systems) and run in a CFX Connect real-time PCR (Biorad).

### Statistical analysis

The data were assessed for normality and equal variance with QQ plots and plotting the residuals, respectively, using the Prism program for statistical analyses (Graphpad). If the data (untransformed or transformed) met the criteria for an ordinary ANOVA, it was performed and multiple pairwise comparison *post hoc* analyses were performed with Tukey’s or Dunnett’s test. If the criteria for an ordinary ANOVA were not met, a non-parametric Kruskal Wallis test (ANOVA of ranks) was performed followed by multiple pairwise comparisons *post hoc* analyses were performed with Dunn’s test.

## Results and Discussion

### The screen for rescue of a hypomorphic dpp wingless phenotype

Flybase was searched with the GO term “receptor ligand activity” in 2022 identifying 106 genes (Öztürk-Çolak et al., 2024). These genes were screened for the availability of a GAL4 UAS construct inserted on the third chromosome in a Drosophila stock center. 45 UAS stocks were identified for 39 genes (Table S1 and S3). 40 of 45 crosses representing 35 genes yielded progeny of the correct genotype to assess rescue (Table S3). The *dpp*^*d5*^*/dpp*^*d6*^ wing phenotype is a strongly reduced wing that still retains the wing costal cell and alula (Figure 1a; Jürgens et al., 2024; for a wild-type wing see Figure 2a). When expressed in a *dpp*^*d5*^*/dpp*^*d6*^ background (Figure 1a) using an imaginal disc specific *dppGAL4* driver, 3 UAS constructs expressing the coding regions derived from 3 genes, *hh, Sema-a1* and *Wnt2* partially rescued the dpp phenotype (Figure 1 b, c, d). Quantification of the male and female wing area of flies expressing either HH, SEMA1A or WNT2 verified the visual partial rescue with expression of any of the 3 in females but only WNT2 in males (Figure 1 e). The rescue by HH is very variable in males and females with some strong increases in wing area in females (*P*=0.008) and with some males strongly rescued, but overall not found significant; however, expression of SEMA-A1 rescued wing growth in females (*P*=0.001) but not males (*P*=1) indicating sexually dimorphic rescue. About a third of the 106 genes identified in 2022 encoding receptor ligand activity were screened and 3 rescues of the dpp hypomorphic phenotype were identified suggesting that screening all 106 genes may identify 4-8 additional examples of partial rescue and potentially an example(s) of strong rescue. Repeating the search of Flybase with the GO term “receptor ligand activity” in 2026 increased the number of genes identified to 118 adding to the potential of identifying additional examples of rescue. The high frequency of rescue observed in this screen suggests that screens for rescues of other phenotypes due to the loss of expression of other SLs have a more than reasonable chance of successfully identifying phenotypic nonspecificity as was found in repeated screens for rescue of 8 of 9 distinct TF phenotypes (Percival-Smith, 2017; Percival-Smith et al., 2023).

**Figure 1.**
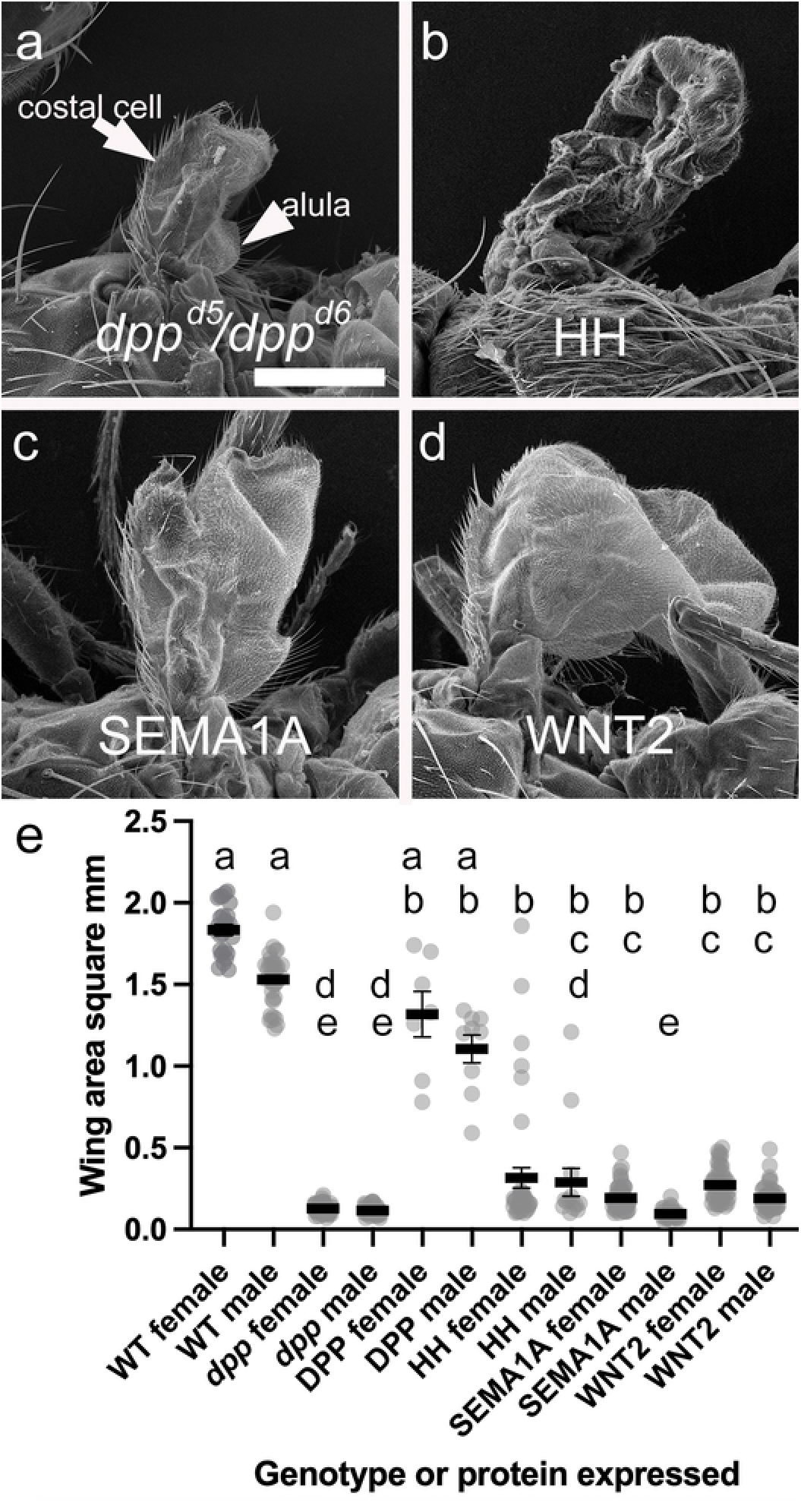
Rescue of the *dpp*^*d5*^*/dpp*^*d6*^ hypomorphic wing phenotype by the expression of HH, SEMA-A1 or WNT2. Panels a-d are scanning electron micrographs of adult flies that are either a *y w; dpp*^*d5*^*/dpp*^*d6*^; *dppGAL4*,b *y w; dpp*^*d5*^*/dpp*^*d6*^; *dppGAL4/P*{*UAShh, w*^+^*}*,c *y w; dpp*^*d5*^*/dpp*^*d6*^; *dppGAL4/M*{*UASSema-a1, w*^+^*}*, or d *y w; dpp*^*d5*^*/dpp*^*d6*^; *dppGAL4/M*{*UASWnt2, w*^+^*}*. The scale bar in panel a is 200 μm and is same for panels b-d. The costal cell of the wing is indicated by an arrow and the alula is indicated by an arrowhead in panel a. Panel e is a scatter plot of wing area of male and female flies. The mean and SD are indicated. Analysis of the data with an ANOVA of ranks detected differences (H(12) = 325, *P*<0.0001) and *post hoc* multiple pair-wise comparisons were performed with Dunn’s test. The data that are not different (*P*>0.05) have the same letter. WT wild type; dpp *y w; dpp*^*d5*^*/dpp*^*d6*^; *dppGAL4*; DPP *y w; dpp*^*d5*^*/dpp*^*d6*^; *dppGAL4/P*{*UASdpp, w+}*; HH *y w; dpp*^*d5*^*/dpp*^*d6*^; *dppGAL4/P*{*UAShh, w*^+^*}*; SEMA-A1 *y w; dpp*^*d5*^*/dpp*^*d6*^; *dppGAL4/M*{*UASsema-a1, w*^+^*}*; and WNT2 *y w; dpp*^*d5*^*/dpp*^*d6*^; *dppGAL4/M*{*UASWnt2, w*^+^*}*. Panels a-d are scanning electron micrographs showing the rescue of wing growth b-d relative to the mutant control a. Panel e is a scatter plot of the quantification of wing areas.

**Figure 2.**
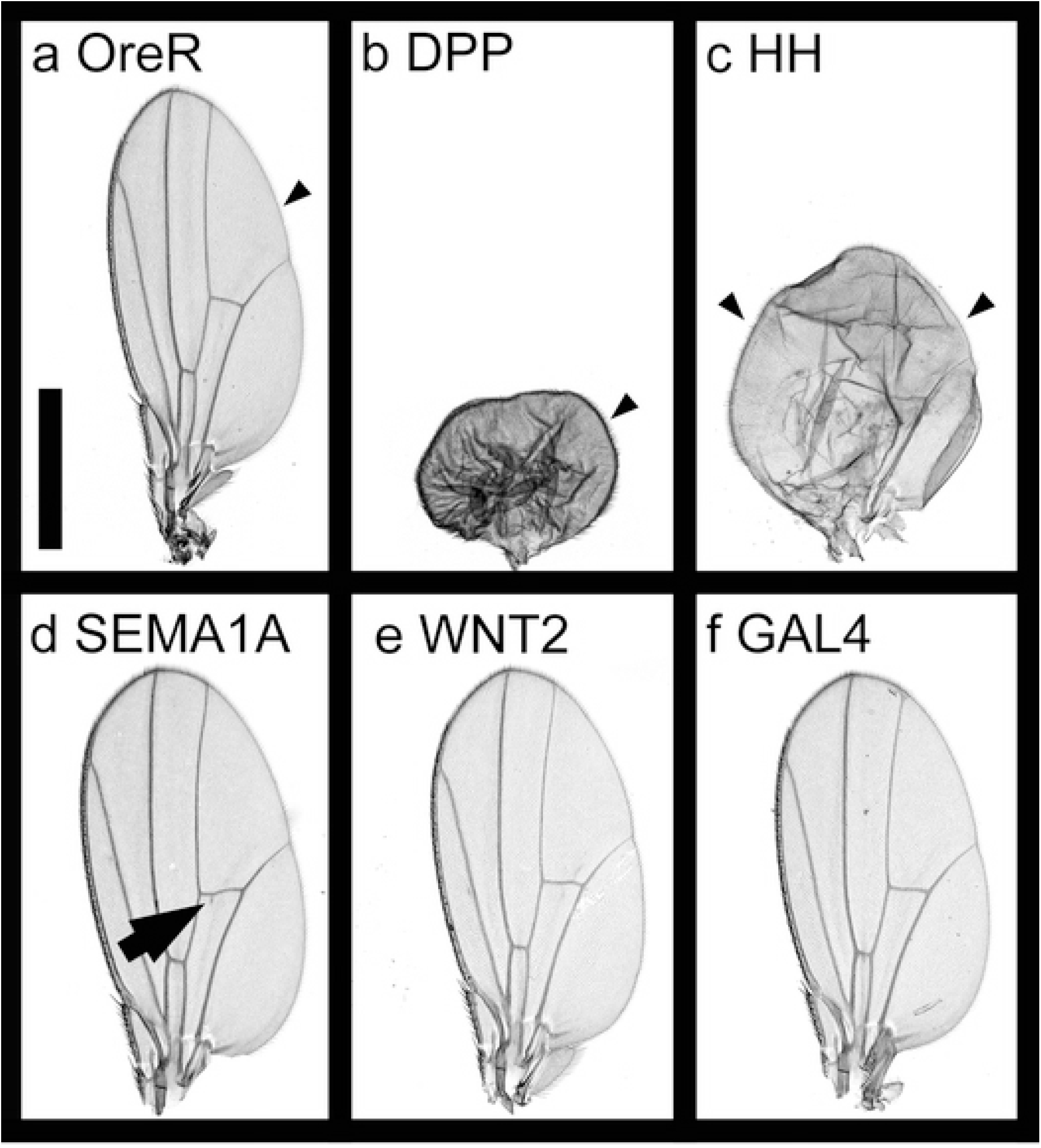
Ectopic expression of DPP, HH, SEMA1A and WNT2 in the wing disc with the *nubbinGAL4* driver. All panels are female wings. a OreR wild type flies; b expressing DPP; c expressing HH; d expressing SEMA1A; e expressing WNT2; f expressing GAL4 from *nubGAL4* alone. The bar in panel a indicates 500μm. The arrowheads in panels a-c indicate the row of tricombs characteristic of the posterior wing compartment, and the arrow in panel d indicates the vein spur on the posterior crossvein. Panels a-f are brightfield images of wings. a and f are control wild-type a and expressing GAL4 only f. Panels b-e express DPP, HH, SEMA1A and WNT2, respectively, with the *nubGAL4* driver.

### Activation of DPP expression as a mechanism for rescue of the hypomorphic dpp phenotype

The role of HH in activating DPP expression required for wing growth and pattern formation is well described (Tabata and Kornberg 1994) and is a potential explanation for the partial rescue of the dpp hypomorphic phenotype. Overexpression of HH with the *dppGAL4* driver may increase expression of *DPP* mRNA from the two hemizygous hypomorphic allelic loci resulting in increased DPP signalling activity and partial rescue. Therefore, raising the question of whether WNT2 or SEMA1A also activate DPP expression like HH rather than substituting for DPP. Due to the embryonic lethality of *dpp* null alleles, this question cannot be addressed directly with an analysis of genetic epistasis in adult wing growth. Instead, a gain-of-function phenotype induced by ectopic expression of HH in the wing imaginal disc with the driver *nubbinGAL4* was assessed. Ectopic expression of HH throughout the wing pouch with the *nubGAL4* driver resulted in a circular wing with the anterior compartment transformed to posterior identity; the anterior wing margin lost its characteristic rows of bristles and was replaced with a tricomb margin characteristic of the posterior wing margin (Figure 2c). In addition, vein formation was reduced. Ectopic expression of DPP with *nubbinGAL4* resulted in a small, circular wings with no vein formation, but the anterior compartment was not transformed to posterior identity. The inability of the DPP wing to inflate post-eclosion explains its smaller size relative to the HH wing (Figure 2 b). The ectopic expression of HH and DPP induce similar phenotypes as expected from them being on the same developmental pathway. Ectopic expression of both SEMA1A and WNT2 did not affect wing formation like ectopic expression of either HH or DPP. WNT2 wings were wild-type (Figure 2e, a); while SEMA1A wings had a small vein spur off the posterior crossvein (Figure 2d; Jürgens et al., 2024). The ectopic expressions with the *nubbinGAL4* driver do not support an obvious role for WNT2 and SEMA1A, like HH, in activating DPP expression as a mechanism for partial rescue.

### WNT2/Frizzled 4 (FZ4) and SEMA1A/Plexin A (PLEXA) requirement in wing formation

The rescue of the hypomorphic dpp phenotype by SEMA1A and WNT2 non-resident SLs may occur by the non-resident SL either having a role along the DPP pathway downstream of DPP, or by the ectopic expression of the SL complementing the loss of DPP expression. If the non-resident SL has a role in the DPP pathway, the SL will be required for wing growth, and if the SL complements the DPP pathway, it will not be required for wing growth.

Both *WNT2* and *SEMA1A* mRNA accumulate in wing imaginal discs (Table S4; Sidhu and Percival-Smith, 2025). *WNT2* mRNA accumulates in the wing pouch (Figure 3d; Everetts et al., 2021; Yu et al., 2020). To assess the role of WNT2 in wing development, the wing area of *Wnt2*^*O*^*/Wnt2*^*KO*^ null hemizygotic females were measured, and a size decrease was not detected suggesting that WNT2 does not promote wing growth (Figure 3j; Kozopas et al., 1998; Yu et al., 2020). Rather, the small increase in wing area suggests that WNT2 is required for suppression/restriction of wing growth. Therefore, WNT2 activity is not downstream/epistatic to DPP in the DPP dependent wing growth pathway suggesting that WNT2 substitutes for the loss of DPP. However, it is puzzling how ectopic and likely overexpression of WNT2 with *dppGAL4* results in partial rescue of wing growth when WNT2 is already expressed in the wing pouch and required for restriction of wing growth.

**Figure 3.**
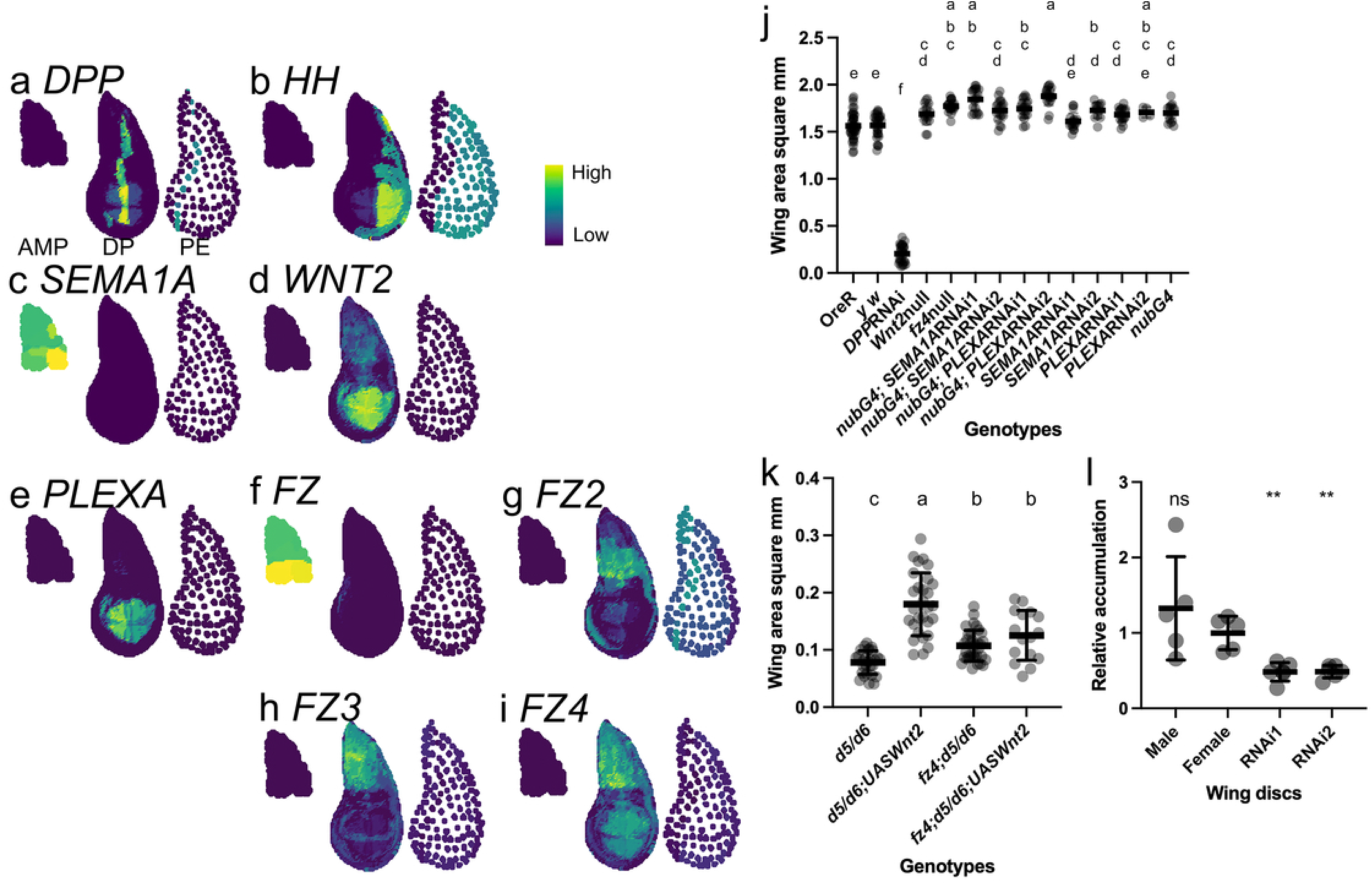
The expression and requirement of SEMA1A and WNT2 for wing growth. Panels a-i are virtual insitus generated using the data and method of Everetts et al., 2021. Each panel from left to right shows the cells of the adult muscle precursor cells (AMP), the disc proper (DP) and the peripodial envelop (PE). Panel a *DPP* mRNA accumulation; b *HH* accumulation; c *SEMA1A* accumulation; d *WNT2* accumulation; e *PLEXA* accumulation; f *FZ* accumulation; g *FZ2* accumulation; h *FZ3* accumulation; and i *FZ4* accumulation. The mRNA accumulation scale in shown to the right of panel b. Panel j is a scatter plot of the wing areas of female wings assessing the requirements of SEMA1A and WNT2 for wing growth. The wing area is on the y-axis and the genotypes are indicated below the x-axis. The data was analyzed with an ordinary ANOVA (F_13, 276_=441; *P*<0.0001). Data with the same letters are not different (*P*>0.05). *SEMA1A RNAi1* 29554; *SEMA1ARNAi2* 34320; *PLEXARNAi1* 30483; and *PLEXARNAi2* 67845 (Table S1). Panel k is a scatter plot of the wing area of female wings assessing the requirement for FZ4 for WNT2 dependent wing growth. The wing area is on the y-axis and the genotypes are indicated below the x-axis. The data was analyzed with an ordinary ANOVA of log transformed data (F_3, 103_=37; *P*<0.0001). Data with the same letter are not different (*P*>0.05). Panel l is a scatter plot of relative *PLEXA* mRNA accumulation. The accumulations are relative to the average accumulation in wild type female wing imaginal discs. The data was analyzed with an ordinary ANOVA of log transformed data (F_3, 18_=13; *P*=0.0001). The ** indicate *P*<0.01 relative to *PLEXA* accumulation in female wing imaginal discs; ns not significant (*P*>0.05). The mean and SD are indicated in panels j-l. Panels a-i are virtual insitus showing the accumulation of the transcripts *DPP, HH, SEMA1A, WNT2, PLEXA, FZ, FZ2, FZ3*, and *FZ4*, respectively. Panel j is a scatter plot of wing area showing the SEMA1A and WNT2 are not required for growth. Panel k is a scatter plot of wing area showing that FZ4 is required for WNT2 partial rescue. Panel l is a scatter plot of *PLEXA* mRNA accumulation in males, females, and when knocked down with RNAi.

All the Frizzled WNT receptors are expressed in the wing imaginal disc (Table S4; Figure 3f-i) with *FZ4* mRNA accumulating strongly in the wing pouch (Figure 3i). Inclusion of a *fz4*^*3-1*^ null allele in flies expressing WNT2 in a *dpp*^*d5*^*/dpp*^*d6*^ mutant suppressed WNT2 dependent partial wing growth rescue indicating that FZ4 is an important WNT2 receptor in the wing (Figure 3k; McElwain et al., 2011). Both FZ4 and WNT2 restrict wing area (Figure 3j) and *fz4; dpp*^*d5*^*/dpp*^*d6*^ wings were larger than *dpp*^*d5*^*/dpp*^*d6*^ wings confirming the requirement of WNT2/FZ4 in suppression of wing growth (Figure 3k). The requirement for suppression of wing growth by FZ4 in *dpp*^*d5*^*/dpp*^*d6*^ wings suggests that WNT2 expression is present in *dpp*^*d5*^*/dpp*^*d6*^ developing wing discs. This observation does not provide support for the proposal that DPP expression activates WNT2 expression to serve as a negative feedback factor in suppression of wing growth. Importantly, the requirement of FZ4 indicates that WNT2 does not stimulate DPP dependent wing growth by weakly binding to the Thickvein Punt receptor and weakly activating the DPP pathway.

*SEMA1A* mRNA accumulates in the adult muscle precursor (AMP) cells of wing imaginal discs (Figure 3c; Everetts et al., 2021). Although *SEMA1A* mRNA accumulates preferentially in AMP cells, nonetheless, its requirement for wing growth was assessed in the wing pouch. Independent RNAi hairpins that target *SEMA1A* mRNA and Dicer2 (DCR2) to increase the effectiveness of RNAi were co-expressed with a *nubbinGAL4* driver in the wing pouch (Dietzl et al., 2007). No requirement of SEM1A for wing growth was found (Figure 3j). However, this experiment unlikely affected accumulation of *SEMA1A* in the AMP cells but, interestingly, the SEMA1A receptor, *PLEXA*, mRNA is expressed in the wing pouch (Figure 3e; Winberg et al., 1998). The accumulation of *PLEXA* mRNA in the wing disc was reduced 50% with RNAi expression expressed with *nubbinGAL4* from two independent RNAis (Figure 3l). In flies co-expressing DCR2 with the RNAis to increase the effectiveness of RNAi, no reduction in the wing area was observed (Dietzl et al., 2007). The observed increase in wing area in all the knockdown wings is also observed in all the *UASRNAi* stocks and with expression of GAL4 from *nubbinGAL4* alone indicating that the increase in wing area is likely due to the genetic background and not a requirement for wing growth suppression (Figure 3j). *PLEXA* mRNA accumulated in both male and female wing imaginal discs ruling out sex-specific accumulation of *PLEXA* mRNA as the explanation for SEMA1A, sex-specific rescue (Figure 3l). Flies homozygous for loss-of-function *Sema1a* alleles are partially adult viable, develop wings but are flightless (Kolodkin et al., 1993). These knockdown experiments, accumulation of *SEMA1A* mRNA and previous analysis of *Sema1a* loss-of-function alleles (Kolodkin et al., 1993) provide no evidence for a role of *SEMA1A/PLEXA* mRNA in wing growth. In summary, expression of WNT2/FZ4 and SEMA1A/PLEXA are not required for wing growth, and therefore, the mechanism of rescue is likely substitution of DPP by WNT2 and SEMA1A.

## Conclusions and future studies

The expression of DPP, HH, SEMA1A and WNT2 rescue a hypomorphic dpp adult wing phenotype and the expression of SEMA1A and WNT2 likely substitute for DPP. These observations are an expectation of the functional redundancy hypothesis of signal transduction terminal TF function. In addition, that phenotypic nonspecificity is observed in 8 of 9 *TF* loci tested so far suggests that phenotypic nonspecificity of SL function will be observed in screens of other *SL* loci (Percival-Smith et al., 2023). Phenotypic nonspecificity is an evolutionary opportunity; whereas, phenotypic specificity is an evolutionary constraint (Percival-Smith 2018). Phenotypic nonspecificity allows the replacement of one developmental pathway with another during evolution (Percival-Smith et al., 2023). Phenotypic nonspecificity may be important for how a pathway generally associated with immunity in animals has become associated with dorsoventral patterning in many insect orders (Pechmann et al., 2021). With the observations of phenotypic nonspecificity of both TF and SL function, there is no reason to assume phenotypic specificity for the requirements of kinases, phosphatases and RNA binding proteins. The null hypothesis of the assumption of phenotypic specificity is easily tested in screens for phenotypic nonspecificity.

## Data Availability Statement

All fly stocks were obtained from either the Bloomington or FlyORF stock centers (Table S1).

## Acknowledgements

We thank the Bloomington and FlyORF Drosophila stock centers for providing the various fly stocks used in this study. We thank Karen Nygard and Ivan Barker for assistance with microscopy and SEM, respectively. The wing areas and critical point drying were performed in the Biotron Integrated Microscopy Laboratory and the SEM images collected at Surface Science Western (London ON). We thank Gurpreet Dhami for assistance with qPCR and Gabriella Sidhu for generating the virtual insitu images.

## Contributions

AP-S conceived the study. CB searched for available UAS insertions. CB and AP-S screened for rescue. AP-S collected SEM images, quantified wing area, assessed requirement of SEMA1A and WNT2, performed qPCR, assessed HH phenocopy and assessed the requirement of FZ4 for WNT2 signalling. AP-S wrote the first draft of the manuscript and both AP-S and CB revised the draft.

